# Not one size fits all: influence of EEG type when training a deep neural network for interictal epileptiform discharge detection

**DOI:** 10.1101/2023.03.02.530752

**Authors:** Catarina da Silva Lourenço, Marleen C. Tjepkema-Cloostermans, Michel J. A. M. van Putten

## Abstract

**Objective:** Deep learning methods have shown potential in automating interictal epileptiform discharge (IED) detection in electroencephalograms (EEGs). While it is known that these algorithms are dependent on the type of data used for training, this has not been explored in EEG analysis applications. We study the difference in performance of deep learning algorithms on routine and ambulatory EEG data.

**Methods:** We trained the same neural network on three datasets: 166 routine EEGs (VGGC–R), 75 ambulatory EEGs (VGGC–A) and a combination of the two data types (VGGC-C, 241 EEGs total). Networks were tested on 34 routine EEGs and 33 ambulatory recordings, where all 2 s non-overlapping epochs were labeled with a probability that expressed the likelihood of containing an epileptiform discharge. Performance was quantified as sensitivity, specificity and the rate of false detections (FPR).

**Results:** The VGGC-R led to 84% sensitivity at 99% specificity on the routine EEGs, but its sensitivity was only 53% on ambulatory EEGs, with a FPR > 3 FP/min. The VGGC-C and VGGC-A yielded sensitivities of 79% and 60%, respectively, at 99% specificity on ambulatory data, but their sensitivity was less than 60% for routine EEGs.

**Conclusion:** We show that performance of deep nets for IED detection depends critically on the type of recording. The VGGC-R should be used for routine recordings and the VGGC-C should be used for ambulatory recordings for IED detection.

**Significance:** The type of data used to train algorithms should be optimized according to their application, as this has a significant impact on algorithm performance.

## Introduction

Interictal epileptiform discharges (IEDs) in electroencephalograms (EEGs) reflect an increased likelihood of seizures and their characteristics can assist in the identification and classification of epilepsy syndromes [1, 2]. Visual analysis of EEG recordings by experts is currently the gold standard in IED detection, and a crucial part of epilepsy diagnosis, as well as monitoring the effect of therapy [3].

Visual analysis is a time and resource-consuming endeavor, as EEG review times are long and trained experts are required. Aside from the long learning curve, inter- and intra-subject variability constitute further drawbacks of visual analysis [4, 5]. Given the increasing number of ambulatory and even ultra-long EEG recordings of months to years, made possible by technological developments, the resources needed for this task have increased exponentially [6-8].

Automating IED detection could reduce the number of work hours invested in visual analysis, streamlining procedures in the clinic [2, 9]. This is particularly useful when a large volume of data needs to be analyzed. There have been several approaches aiming to automate IED detection [10]. These range from thresholding of morphological or frequency features [11, 12] to traditional machine learning methods [13, 14] and, more recently, deep learning techniques [15-20]. The popularity of deep learning methods in the medical field has grown in the past years, with applications from predicting mortality from echocardiographic video [21] to skin cancer classification [22]. In EEG analysis, and specifically in epilepsy, deep learning has been applied to tasks such as IED detection, seizure detection and seizure prediction [10, 23-25].

The type and quality of the data used for training and validation of deep learning algorithms is critical, as it also defines the boundary conditions for their applicability. As neural networks learn from experience, they will inevitably make mistakes when exposed to types of data that were not included in the training process [26]. This has been explored in fields such as computer vision or natural language processing, where strategies such as transfer learning are employed to extend model applicability in new contexts [27, 28]. This limitation may also apply to deep learning approaches for IED detection, but this has been left unexplored in current EEG literature. Most studies discussing deep networks for IED detection use routine EEG recordings for training, with an average duration of 20-30 minutes [17-20], while in other works data length and acquisition conditions are not clearly specified [16].

Neural networks trained on EEG data recorded in a hospital environment may perform differently if applied to EEGs recorded in other situations. Routine data is acquired in a more controlled environment and under expert supervision, resulting in less artefacts. Ambulatory data usually contains more and more diverse artefacts, ranging from loose electrodes to electrical interference, chewing, movement, among others. Furthermore, ambulatory EEGs contain periods of sleep, which are not typically present in routine recordings. This implies that rhythms and patterns typical of sleep, such as sleep spindles, K-complexes and vertex waves are widely present in ambulatory recordings but not in routine EEGs [29, 30]. It is unclear whether the performance of a neural network trained with routine EEG data is maintained when applied to ambulatory recordings: while the specific abnormalities (interictal discharges) that need to be detected are essentially similar, performance could be affected by the presence of the other rhythms, transients and artefacts. This is clinically highly relevant, as such recordings are increasingly available and used, among others, for IED detection [6-8].

To the best of our knowledge, there have been no studies comparing the performance of algorithms for IED detection in different types of EEG data. Here, we compare the performance of three deep neural networks trained with different types of EEG data on a test set comprised of routine and ambulatory EEG recordings. We previously reported the performance of the network trained on routine data on an independent test set [15]. The two other algorithms have not been previously published. Further, the test sets used on this study are different and do not overlap with the data used in [15].

## Methods

### EEG data and pre-processing

We used data from 308 patients, ranging from 8 to 86 years old, randomly selected from the database of the Medisch Spectrum Twente, in the Netherlands. Recordings from 159 epilepsy patients were included, as well as 149 EEGs that were classified as normal. In the recordings from epilepsy patients, all the IEDs were visually labeled by experts (MvP and MTC). As EEG is part of routine care, the Medical Ethical Committee Twente waived the need for informed consent. All EEGs were anonymized before analysis.

Of the 308 recordings, 200 were routine EEGs (20±10 minutes) and 108 were ambulatory EEGs (20±1 hours). Routine and ambulatory recordings have significant differences, as ambulatory EEGs include e.g. periods of sleep, chewing, and other artefacts such as electrodes with poor contact, among others, as shown in Figure 1.

**Figure 1.**
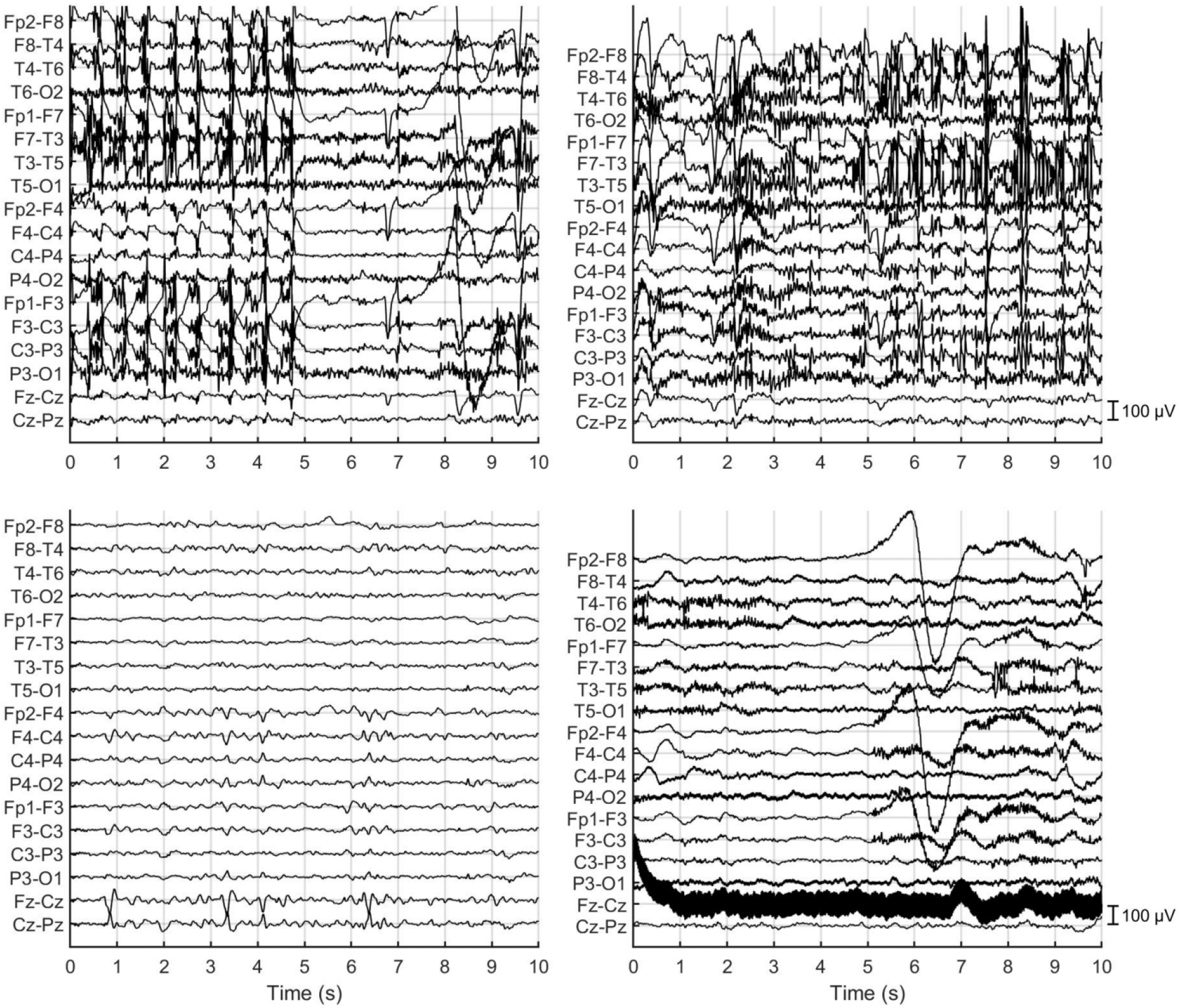
Examples of EEG epochs from ambulatory recordings. The top panels show two chewing artefacts, the bottom left panel includes vertex waves (clearly visible in channels Fz-Cz and Cz-Pz) and the bottom right panel shows an example of an electrode with poor contact (Fz, visible in Fz-Cz). Another artefact in this recording is the large deflection arising from Fp2. Epoch length is 10 s, bipolar montage, filter settings 0.5-30 Hz.

EEG data was filtered in the 0.5–30 Hz range and downsampled to 125 Hz, aiming to reduce artefacts and data dimensionality. The signals, in the longitudinal bipolar montage, were split into 2 s epochs. These steps were implemented in Matlab R2021b (The MathWorks, Inc., Natick, MA).

### Dataset Creation

The EEG data was first separated into independent training and test sets, ensuring that all data epochs from a particular patient were used either for training or for testing. Training data was further divided into three datasets: set R, containing only the routine data; set A, comprised of the ambulatory data and set C, a combination of the previous two datasets. Dataset R had EEGs from 166 subjects, while dataset A contained recordings from 75 subjects. The combination set, C, had data from 241 patients. This is summarized in Table 1.

**Table 1.** Overview of the three training datasets: R, comprised of routine data; A, comprised of ambulatory EEGs and C, including both routine and ambulatory recordings. The number of normal EEGs and EEGs with interictal discharges is shown. The routine training set contained 50 EEGs with focal IEDs and 49 EEGs with generalized IEDs (99 EEGs with IEDs). The ambulatory training set contained 16 EEGs with focal IEDs and 10 EEGs with generalized IEDs (26 EEGs with IEDs). The number of IEDs before and after augmentation is also shown. Augmentation included re-montaging in three different montages and shifting the data acquisition window by 0.5 seconds, resulting in a multiplication of approximately 12 times. It was not an exact 12-fold increase due to some epochs coinciding with end of file, for example.

|  | R | A | C |
| --- | --- | --- | --- |
| # normal EEGs | 67 | 49 | 116 |
| # EEGs with IEDs | 99 | 26 | 125 |
| # IEDs | 2220 | 3262 | 5482 |
| # IEDs after augmentation | 25302 | 37878 | 63180 |

Training data was augmented to increase the number of training samples, as described in [15]. In short: we used three different EEG montages: longitudinal bipolar (DB), Laplacian (SD) and common average (G19). In the SD and G19 montages, the last channel was removed to maintain 18 channels across all samples. Further, to increase the number of IEDs in the dataset, the epochs that contained an IED were shifted with 0 s, 0.5 s, 1 s and 1.5 s. This resulted in the multiplication of the number of IEDs in the datasets by approximately 12 fold (see Table 1), and has shown to result in a significant improvement of performance [15]. As some IEDs could have been overlooked by the expert, we did not use negative samples (i.e. without IEDs) from EEGs with IEDs for training.

Since the test set should be representative of all types of data, we randomly selected 18 routine EEGs with IEDs and 16 routine EEGs that were classified as normal, as well as 16 ambulatory EEGs with IEDs and 17 ambulatory EEGs that were classified as normal (see Table 2). The test set included over 2.300 IEDs (403 from routine EEGs and 1918 from ambulatory EEGs) and 255 h of EEG data. All the samples from all the EEGs (Table 2) were used for testing, and no data augmentation techniques were applied to this dataset.

**Table 2.**
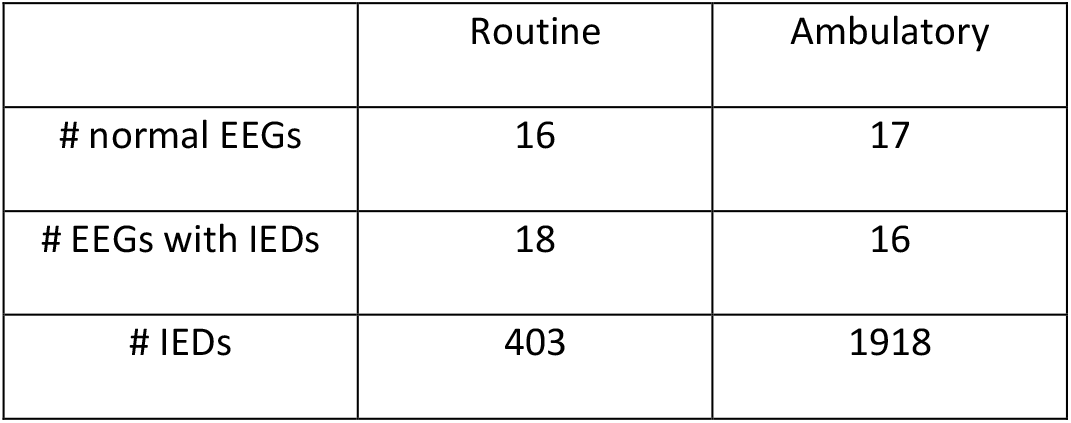
Overview of routine and ambulatory test datasets, including normal EEGs and EEGs with IEDs, including focal IEDs (in 9 routine EEGs and 8 ambulatory EEGs) and generalized IEDs (in 9 routine EEGs and 8 ambulatory EEGs). The number of interictal discharges in the recordings is also shown.

|  | Routine | Ambulatory |
| --- | --- | --- |
| # normal EEGs | 16 | 17 |
| # EEGs with IEDs | 18 | 16 |
| # IEDs | 403 | 1918 |

### Deep Learning models

A modified VGG C [31] convolutional neural network was implemented in Python 3.8 using Tensorflow 2.2.0 and a CUDA–enabled NVIDIA GPU (GTX–1080), running on Rocky Linux 8. The modifications of the original architecture were mostly related to data dimensionality: the input dimensions were changed to [number of epochs x 250 x 18], with 250 corresponding to 2 seconds of data sampled at 125 Hz and 18 being the number of channels. In the final dense layer, we used two nodes instead of 1000 to fit the binary classification problem.

Stochastic optimization was performed using an Adam optimizer [32] with a learning rate of 2^*^10^-5^, β_1_=0.91, β_2_=0.999 and ε=10-8. A sparse categorical cross entropy function was employed to estimate the loss and a batch size of 64 was used.

This architecture was trained with the three datasets: R, A and C, as described in the previous section. This originated the three trained networks: VGGC–R, VGGC–A and VGGC–C. Given the class imbalance of the dataset (the number of samples with IEDs - positive class - was much lower than that of normal samples – negative class), class weights were used. For the VGGC-R (routine model), we used weights of 100:1 (100 corresponding to the positive class, i.e. samples with IEDs) to ensure that the positive class had 100 times more importance for the network when compared to a sample from a healthy control. For the VGGC-A (ambulatory model) and VGGC-C (combined model), the weights were 25:1 and 50:1, respectively. Five-fold cross-validation was used when training the networks with the three different datasets.

### Performance Evaluation

The three neural networks were applied to the test set, classifying each 2 second EEG sample with a probability ranging from 0 (no IED) to 1 (IED is present). The Receiver Operating Characteristic (ROC) curves were plotted for each model, for the routine and ambulatory sets. A threshold of 0.99 was applied to the probabilities, with all EEG samples above this probability value being classified as containing an IED. This threshold was chosen based on optimization done on the training set of each network separately, during cross-validation. At this probability threshold we calculated the sensitivity, specificity and false positive rate (FPR) per minute, including their 95% confidence intervals, for all three networks. An overview of the methods is shown in Figure 2.

**Figure 2.**
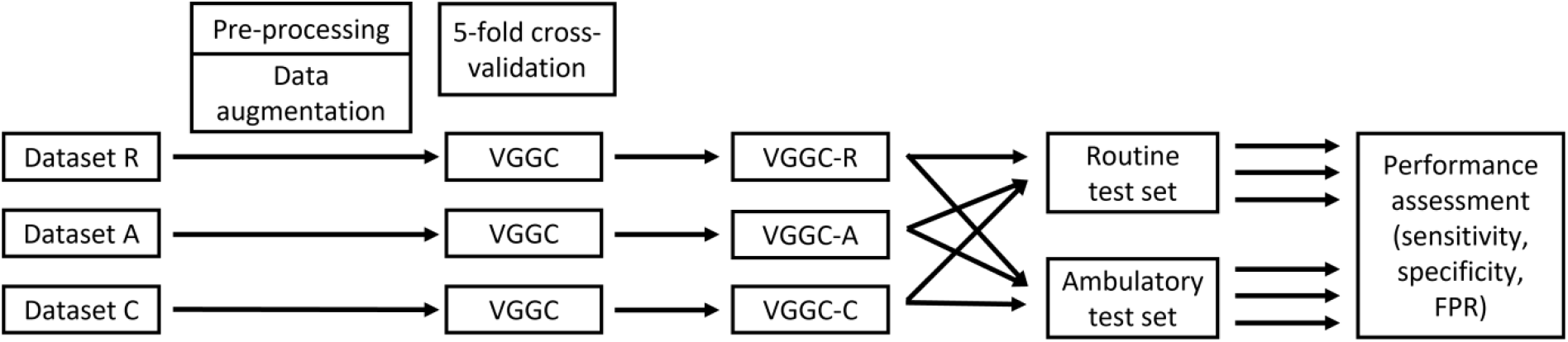
Pipeline of the methodology used in the study. The three datasets described in Table 1 (R, A and C) were pre-processed (filtering, downsampling, splitting into 2 s samples) and augmented (time-shifting and re-referencing to three montages). The VGGC architecture was trained with the three datasets, using 5-fold cross-validation. We named the trained networks VGGC-R, VGGC-A and VGGC-C. These networks processed the EEGs in the routine and ambulatory test sets; performance was assessed with sensitivity, specificity and false positive rate (FPR).

## Results

The classification performance of the three networks (VGGC-R, VGGC-A and VGGC-C) on a routine and on an ambulatory EEG test set is shown in Figure 3. For this study, we focused on the point in the ROC curve corresponding to a probability threshold of 0.99, displayed in Figure 3 as the blue dot and square for the routine and ambulatory EEGs, respectively.

**Figure 3.**
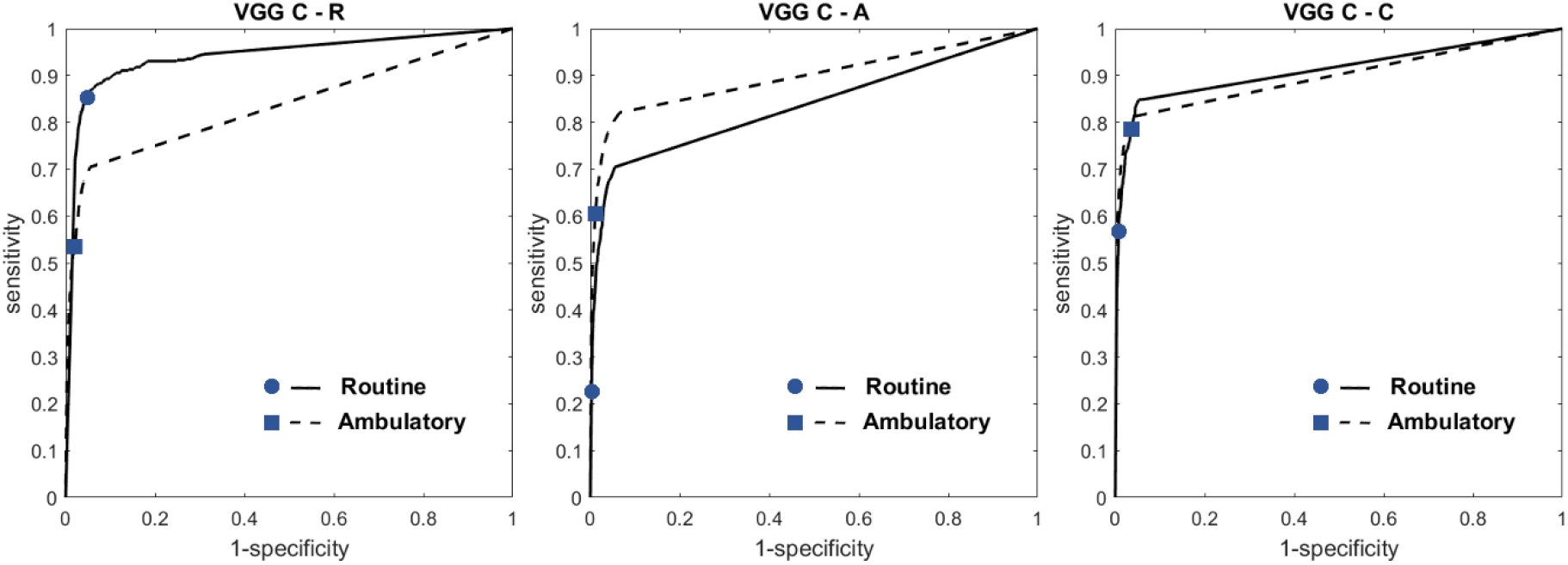
Receiver Operating Characteristic (ROC) curves of the neural networks on the test set with routine and ambulatory EEG data. The left panel shows the VGGC-R, trained on routine data, the middle panel shows the VGGC-A, trained on ambulatory data and the right panel shows the VGGC-C, trained on routine and ambulatory data. The blue dots show the point corresponding to a probability threshold of 0.99 on the routine EEGs and the blue squares show the point corresponding to the same threshold on the ambulatory EEGs. The threshold was optimized in the cross-validation process for each network.

The sensitivity, specificity and FPR at this threshold are shown in Figure 4. For the routine data, the VGGC-R model showed the highest sensitivity, yielding 84% sensitivity at 99% specificity, with a false positive rate of 0.8 FPs/min on these recordings (cf Figure 4). For the ambulatory data, the VGGC-C led to the highest sensitivity of 79% at 99% specificity, with a false positive rate of 0.4 FPs/min (cf Figure 4).

**Figure 4.**
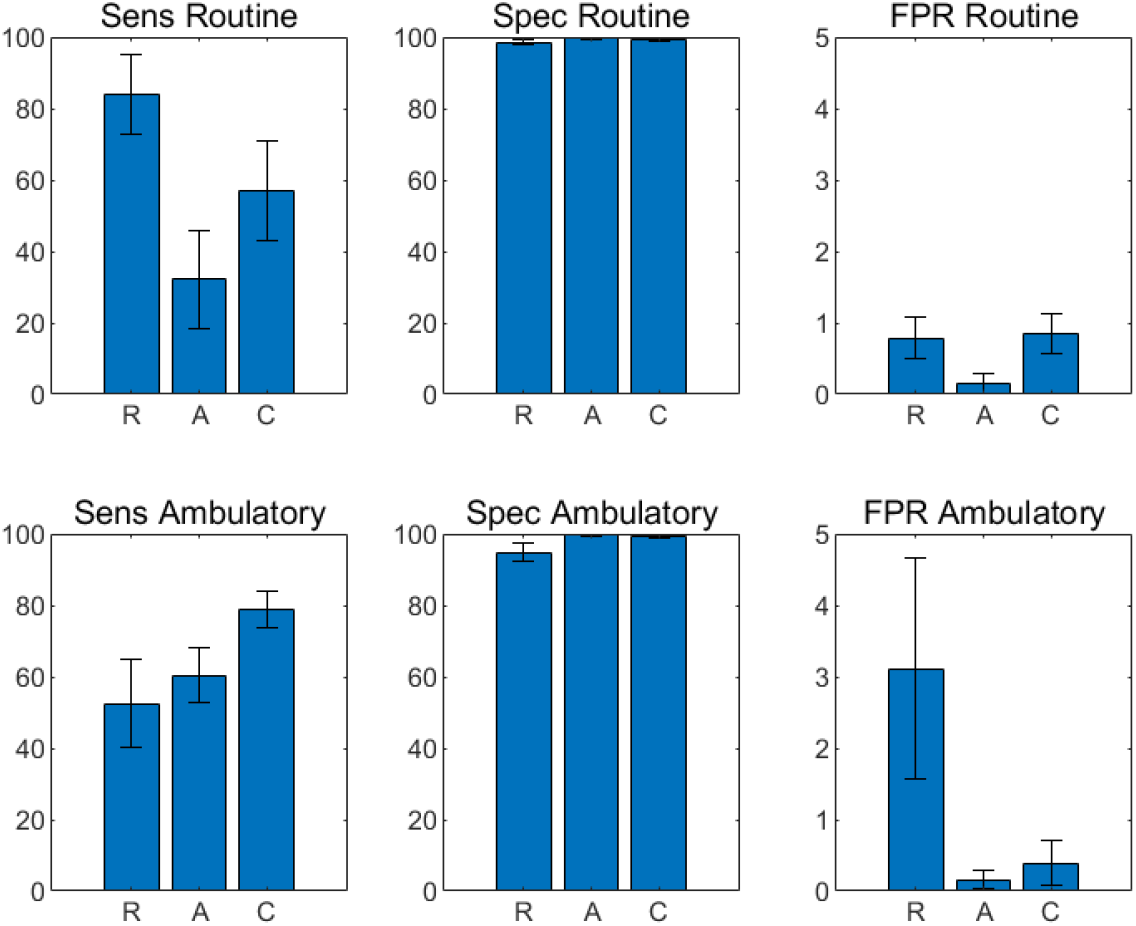
Performance of the of the VGGC-R (R), VGGC-A (A) and VGGC-C (C), thresholded at 0.99, on the routine test set (top) and ambulatory test set (bottom). The panels on the left show the sensitivity (Sens) of the networks, the middle panels report the specificity (Spec) and the panels on the right side illustrate the false positive rate (FPR), in false positives / minute. The 95% confidence intervals are also shown. There is a statistically significant difference (non-overlapping 95% confidence intervals, i.e. p<0.05) between the sensitivity of the VGGC-R and the other two networks on the routine test set. The sensitivity of the VGGC-C is also significantly higher than the sensitivities of the other two networks on the ambulatory test set and the FPR of the VGGC-R is significantly higher than the other two networks on the ambulatory test set.

We also report the sensitivity, specificity and false positive rate of the algorithms on the normal EEGs and on the EEGs with IEDs separately in the Supplementary Material (Tables S1 to S3). The VGGC-R had a FPR of 0.3 FPs/min on normal routine EEGs and 3.2 FPs/min on normal ambulatory EEGs (see Table S1). The FPRs for the VGGC-A and VGGC-C were under 0.8 FPs/min in all the test set sub-partitions (cf Tables S2 and S3). Further, we report the results of the networks for each individual patient (Tables S4 to S9). The VGGC-R and VGGC-C detected IEDs in 17 out of 18 epilepsy patients with routine EEGs and the VGGC-A detected IEDs in 13 out of 18 of the routine EEGs with epileptiform discharges. The three networks detected IEDs in all the ambulatory EEGs containing IEDs.

## Discussion

EEG recordings are essential in the (differential) diagnosis of epilepsy. Routine and ambulatory EEGs have significant differences in duration, but also in the number and type of artefacts and other patterns that can be mistaken as epileptiform. We study the impact of the type of EEG data on the performance of deep neural networks trained for IED detection. As we aim to use these algorithms as a support tool for clinicians, speeding up the IED detection process during EEG review, it is relevant to assess their sensitivity and specificity in different types of EEG data (cf Figures 2 and 3, Tables S1 to S3), as well as their performance on each EEG (Tables S4 to S9).

The three modified VGG C networks show significant differences in sensitivity (cf Figure 4) and overall disparities in performance on the routine and ambulatory test sets (cf Figures 2 and 4), with the best results being achieved by the VGGC-R on the routine test set and by the VGGC-C on the ambulatory test set.

The VGGC-R led to 84% sensitivity at 99% specificity on the routine EEGs and had a particularly low false positive rate on normal routine EEGs (0.3 FPs/min, see Table S1), showing potential for clinical application. In contrast, the sensitivities of the VGGC-A and VGGC-C were under 60%, which is insufficient for clinical use. This is in line with expectations, since the VGGC-R network was trained on routine data, and apparently performed better on data that was acquired in similar conditions as the training dataset.

When tested on the ambulatory recordings, the ROC curves of the VGGC-A and VGGC-C were similar (cf Figure 3). Still, at the 0.99 probability threshold, the VGGC-C showed a significantly higher sensitivity (79% at 99% specificity) for IED detection than the VGGC-A and the VGGC-R (cf Figure 4, Tables S1 to S3). The false positive rate of the VGGC-C on the ambulatory test set (0.4 FPs/min, see Figure 4) was comparable to that of the VGGC-R on the routine test set. The improved performance of the VGGC-C when compared with the VGGC-A in the ambulatory test set likely results from both networks having been exposed to ambulatory data during training, while the VGGC-C ‘saw’, in addition to this, more examples of IEDs from the routine EEG recordings. The larger and more representative number of training samples led to the satisfactory performance of the VGGC-C, with a high sensitivity to IEDs and low sensitivity to artefacts and physiological sleep transients (e.g. K-complexes and vertex waves), making this algorithm potentially applicable to detect IEDs in ambulatory EEGs.

On the ambulatory test set, the VGGC-R performed poorly when compared to the VGGC-A and the VGGC–C as reflected by the low sensitivity (53%, cf Figure 4) and the high false positive rate (> 3 FPs/min for both normal and EEGs with epileptiform discharges, cf Figure 4 and Table S1). This mainly resulted from particular artefacts and sleep phenomena in the ambulatory EEG data, absent in the training set of the VGGC–R, that were erroneously classified as epileptiform discharges.

Most studies concerning automated IED detection used short EEG recordings [17-20]. From these, some algorithms were trained with a low volume of data (e.g. [18], routine EEGs from 5 patients) or used a combination of raw EEGs with additional hand-crafted features for training [19]. One study used EEGs with a median length of 53 minutes to train a convolutional neural network (SpikeNet) [17]. While the authors report only AUC and accuracy, it is possible to infer a false positive rate from the ROC curve. Taking the point where sensitivity=specificity=0.95, the false positive rate is approximately 1.67 FPs/min, assuming 2 s epochs, which is higher than the 0.78 FPs/min from our VGGC-R on the routine dataset. Another study tested an ensemble of one-dimensional convolutional networks on routine EEGs, yielding a sensitivity of 80% with 0.2 FPs/min [20]. This represents a different trade-off between sensitivity and specificity when compared to the VGGC-R, since the sensitivity of our algorithm was larger at the cost of a slightly higher false positive rate.

Other algorithms do not specify the length of the training recordings [16, 25, 33]. DeepSpike [16], a convolutional neural network embedded in the Encevis software, reported a sensitivity of 81.6% at 46.4% specificity for IED detection. The authors used non-epileptiform paroxysmal events instead of normal EEG samples as negative samples for training. This may have contributed to the low specificity of the algorithm, which is subpar to both the VGGC-R on the routine test set and the VGGC-C on the ambulatory test set. Persyst [33], another commercially available software with an IED detection functionality, reported 43.9% sensitivity and 1.65 false detections per minute on 24h recordings. It is not clear whether this algorithm was trained on routine or ambulatory recordings, but the level of performance suggests that it may have been trained on a routine EEG dataset, only. This performance is similar to our VGGC-R tested on the ambulatory dataset, with 52.5% sensitivity and 3.11 FPs/min. In our data, the poor performance results from the mismatch between the type of training and test data, as the VGGC-R yields a much better performance when tested on routine EEGs.

To the best of our knowledge, this is the first work that specifically addresses the importance of the type of EEGs used for training deep learning algorithms for IED detection. Apparently, despite that the interictal discharges generally have similar distinctive features, the presence of other rhythms, non-epileptiform transients (K-complexes, sleep vertex waves) and artefacts in ambulatory recordings affects the performance of networks trained without exposure to these phenomena. Given the large differences in algorithm performance across data types (cf Figure 4, Tables S1 to S3), we argue that networks for IED detection should preferably be applied to similar data as the network used for training.

All of the networks described in this work were trained with EEGs from adults, which is a limitation of our methods as these algorithms cannot be successfully applied to recordings from infants or young children. In this study, we show that algorithm performance depends largely on the type of training data and the patterns that are present in that dataset. Further, training and testing were carried out with data from one clinical center. As patient populations may also vary between hospitals, performance may differ when applied to data from other institutions.

## Conclusion

We show that the type of EEG data has a large impact on the performance of deep learning algorithms trained for IED detection. We suggest that algorithms should be trained with the same type of EEG data they will eventually be applied to, either routine or ambulatory, as this has a significant impact on algorithm performance. In our study, we show that the VGGC-R should be used for routine recordings and the VGGC-C should be used for ambulatory recordings for the best performance in IED detection.

