## Supplementary Material for "Not one size fits all: influence of EEG type when training a deep neural network for interictal epileptiform discharge detection"

*Table S1. Average sensitivity (Sens), specificity (Spec) and false positive rate per minute (FP/min) on the test set for the VGG C network trained with the dataset comprised of routine EEG data (R). The 95% confidence intervals (CI) are shown for each metric. The test data is split into routine EEGs containing IEDs (Epi rout), ambulatory EEGs containing IEDs (Epi amb), normal routine EEGs (N rout) and normal ambulatory EEGs (N amb).*

| VGGC-R | Sens | Spec | FP / min |
|---|---|---|---|
| Epi rout | 84.0 (CI 73.0-95.0) | 96.2 (CI 94.7-97.6) | 1.24 (CI 0.83-1.65) |
| Epi amb | 52.5 (CI 40.2-64.9) | 94.9 (CI 92.4-97.4) | 3.05 (CI 1.54-4.56) |
| N rout | | 99.5 (CI 99.2-99.9) | 0.27 (CI 0.00-0.72) |
| N amb | | 94.7 (CI 90.3-99.2) | 3.16 (CI 0.45-5.87) |

*Table S2. Average sensitivity (Sens), specificity (Spec) and false positive rate per minute (FP/min) on the test set for the VGG C network trained with the dataset comprised of ambulatory EEG data (A). The 95% confidence intervals (CI) are shown for each metric. The test data is split into routine EEGs containing IEDs (Epi rout), ambulatory EEGs containing IEDs (Epi amb), normal routine EEGs (N rout) and normal ambulatory EEGs (N amb).*

| VGGC-A | Sens | Spec | FP / min |
|---|---|---|---|
| Epi rout | 32.2 (CI 18.4-46.0) | 99.5 (CI 99.1-100.0) | 0.27 (CI 0.02-0.51) |
| Epi amb | 60.4 (CI 52.6-68.1) | 99.5 (CI 99.1-99.9) | 0.29 (CI 0.04-0.53) |
| N rout | | 100.0 (CI 99.9-100.0) | 0.02 (CI 0.00-0.11) |
| N amb | | 99.9 (CI 99.0-100.0) | 0.05 (CI 0.00-0.11) |

*Table S3. Average sensitivity (Sens), specificity (Spec) and false positive rate per minute (FP/min) on the test set for the VGG C network trained with the dataset comprised of routine and ambulatory EEG data (C). The 95% confidence intervals (CI) are shown for each metric. The test data is split into routine EEGs containing IEDs (Epi rout), ambulatory EEGs containing IEDs (Epi amb), normal routine EEGs (N rout) and normal ambulatory EEGs (N amb).*

| VGGC-C | Sens | Spec | FP / min |
|---|---|---|---|
| Epi rout | 57.0 (CI 43.0-71.1) | 98.7 (CI 97.8-99.6) | 0.72 (CI 0.22-1.22) |
| Epi amb | 78.8 (CI 73.8-83.9) | 99.0 (CI 98.1-100.0) | 0.57 (CI 0.00-1.16) |
| N rout | | 99.9 (CI 99.8-100.0) | 0.05 (CI 0.00-0.22) |
| N amb | | 99.6 (CI 99.3-100.0) | 0.22 (CI 0.02-0.42) |

*Table S4. Performance of the VGGC–R on the routine test set, comprised of epileptic and normal EEGs. Each line represents an EEG of a test patient. At the probability threshold of 0.99, the number of true positives (TP), false positives (FP), false negatives (FN) and true negatives (TN) are shown.*

| | TP | FP | FN | TN |
|---|---|---|---|---|
| Epileptic | 0 | 1 | 1 | 601 |
| | 14 | 2 | 3 | 1009 |
| | 12 | 3 | 3 | 560 |
| | 12 | 9 | 1 | 569 |
| | 3 | 14 | 0 | 579 |

| | | | | |
|---|---|---|---|---|
| | 19 | 25 | 3 | 510 |
| | 11 | 17 | 1 | 512 |
| | 141 | 20 | 7 | 222 |
| | 12 | 3 | 0 | 455 |
| | 43 | 19 | 0 | 595 |
| | 8 | 20 | 0 | 565 |
| | 17 | 10 | 0 | 546 |
| | 6 | 11 | 3 | 666 |
| | 33 | 24 | 4 | 516 |
| | 4 | 15 | 0 | 718 |
| | 10 | 10 | 5 | 618 |
| | 1 | 8 | 0 | 611 |
| | 16 | 4 | 10 | 1129 |
| Normal | 0 | 0 | 0 | 598 |
| | 0 | 11 | 0 | 646 |
| | 0 | 17 | 0 | 593 |
| | 0 | 0 | 0 | 600 |
| | 0 | 0 | 0 | 565 |
| | 0 | 1 | 0 | 621 |
| | 0 | 1 | 0 | 614 |
| | 0 | 0 | 0 | 604 |
| | 0 | 1 | 0 | 629 |
| | 0 | 0 | 0 | 601 |
| | 0 | 4 | 0 | 636 |
| | 0 | 4 | 0 | 594 |
| | 0 | 5 | 0 | 593 |
| | 0 | 0 | 0 | 603 |
| | 0 | 0 | 0 | 603 |
| | 0 | 2 | 0 | 791 |

*Table S5. Performance of the VGGC–A on the routine test set, comprised of epileptic and normal EEGs. Each line represents an EEG of a test patient. At the probability threshold of 0.99, the number of true positives (TP), false positives (FP), false negatives (FN) and true negatives (TN) are shown.*

| | TP | FP | FN | TN |
|---|---|---|---|---|
| Epileptic | 0 | 0 | 1 | 602 |
| | 0 | 0 | 17 | 1011 |
| | 8 | 0 | 7 | 563 |
| | 8 | 3 | 5 | 575 |
| | 3 | 22 | 0 | 571 |
| | 11 | 0 | 11 | 550 |
| | 6 | 2 | 6 | 577 |
| | 5 | 3 | 143 | 323 |
| | 10 | 1 | 2 | 457 |
| | 15 | 1 | 28 | 613 |
| | 3 | 0 | 5 | 585 |

| | | | | |
|---|---|---|---|---|
| | 0 | 1 | 17 | 555 |
| | 0 | 1 | 9 | 676 |
| | 6 | 11 | 31 | 609 |
| | 2 | 1 | 2 | 762 |
| | 3 | 2 | 12 | 626 |
| | 0 | 0 | 1 | 619 |
| | 5 | 0 | 21 | 1133 |
| Normal | 0 | 0 | 0 | 598 |
| | 0 | 0 | 0 | 657 |
| | 0 | 4 | 0 | 606 |
| | 0 | 0 | 0 | 600 |
| | 0 | 0 | 0 | 656 |
| | 0 | 0 | 0 | 622 |
| | 0 | 0 | 0 | 614 |
| | 0 | 0 | 0 | 604 |
| | 0 | 0 | 0 | 630 |
| | 0 | 0 | 0 | 601 |
| | 0 | 0 | 0 | 640 |
| | 0 | 0 | 0 | 598 |
| | 0 | 0 | 0 | 598 |
| | 0 | 0 | 0 | 603 |
| | 0 | 0 | 0 | 603 |
| | 0 | 0 | 0 | 793 |

*Table S6. Performance of the VGGC–C on the routine test set, comprised of epileptic and normal EEGs. Each line represents an EEG of a test patient. At the probability threshold of 0.99, the number of true positives (TP), false positives (FP), false negatives (FN) and true negatives (TN) are shown.*

| | TP | FP | FN | TN |
|---|---|---|---|---|
| Epileptic | 0 | 0 | 1 | 602 |
| | 2 | 2 | 15 | 1009 |
| | 9 | 0 | 6 | 563 |
| | 10 | 5 | 3 | 573 |
| | 3 | 20 | 0 | 573 |
| | 15 | 3 | 7 | 547 |
| | 8 | 41 | 4 | 538 |
| | 57 | 15 | 91 | 311 |
| | 12 | 3 | 0 | 455 |
| | 39 | 7 | 4 | 607 |
| | 6 | 3 | 2 | 582 |
| | 12 | 0 | 5 | 556 |
| | 2 | 2 | 7 | 675 |
| | 18 | 23 | 19 | 597 |
| | 1 | 2 | 3 | 761 |
| | 5 | 1 | 10 | 627 |
| | 1 | 0 | 0 | 619 |

| | | | | |
|---|---|---|---|---|
| | 10 | 2 | 16 | 1131 |
| Normal | 0 | 0 | 0 | 598 |
| | 0 | 1 | 0 | 656 |
| | 0 | 7 | 0 | 603 |
| | 0 | 0 | 0 | 600 |
| | 0 | 0 | 0 | 656 |
| | 0 | 0 | 0 | 622 |
| | 0 | 0 | 0 | 614 |
| | 0 | 0 | 0 | 604 |
| | 0 | 0 | 0 | 630 |
| | 0 | 0 | 0 | 601 |
| | 0 | 1 | 0 | 639 |
| | 0 | 0 | 0 | 598 |
| | 0 | 0 | 0 | 598 |
| | 0 | 0 | 0 | 603 |
| | 0 | 0 | 0 | 603 |
| | 0 | 0 | 0 | 793 |

*Table S7. Performance of the VGGC–R on the ambulatory test set, comprised of epileptic and normal EEGs. Each line represents an EEG of a test patient. At the probability threshold of 0.99, the number of true positives (TP), false positives (FP), false negatives (FN) and true negatives (TN) are shown.*

| | TP | FP | FN | TN |
|---|---|---|---|---|
| Epileptic | 57 | 394 | 59 | 17938 |
| | 124 | 671 | 43 | 39704 |
| | 125 | 7964 | 36 | 29906 |
| | 3 | 143 | 21 | 37663 |
| | 52 | 1699 | 22 | 37141 |
| | 86 | 3422 | 94 | 29208 |
| | 3 | 1255 | 1 | 39950 |
| | 44 | 1350 | 25 | 38736 |
| | 29 | 228 | 25 | 38406 |
| | 40 | 749 | 74 | 36535 |
| | 539 | 3527 | 33 | 35229 |
| | 32 | 638 | 60 | 33117 |
| | 131 | 2403 | 19 | 34809 |
| | 3 | 152 | 11 | 39185 |
| | 29 | 1354 | 86 | 15005 |
| | 4 | 2002 | 18 | 28526 |
| Normal | 0 | 98 | 0 | 39270 |
| | 0 | 436 | 0 | 38332 |
| | 0 | 1061 | 0 | 36805 |
| | 0 | 99 | 0 | 39377 |
| | 0 | 725 | 0 | 41043 |
| | 0 | 12186 | 0 | 25610 |
| | 0 | 3865 | 0 | 36211 |

| | | | | |
|---|---|---|---|---|
| | 0 | 492 | 0 | 39176 |
| | 0 | 272 | 0 | 38496 |
| | 0 | 156 | 0 | 38310 |
| | 0 | 482 | 0 | 39486 |
| | 0 | 10433 | 0 | 27433 |
| | 0 | 341 | 0 | 31874 |
| | 0 | 297 | 0 | 32034 |
| | 0 | 373 | 0 | 7468 |
| | 0 | 755 | 0 | 31091 |
| | 0 | 398 | 0 | 29723 |

*Table S8. Performance of the VGGC–C on the ambulatory test set, comprised of epileptic and normal EEGs. Each line represents an EEG of a test patient. At the probability threshold of 0.99, the number of true positives (TP), false positives (FP), false negatives (FN) and true negatives (TN) are shown.*

| | TP | FP | FN | TN |
|---|---|---|---|---|
| Epileptic | 60 | 44 | 56 | 18288 |
| | 85 | 91 | 82 | 40284 |
| | 72 | 1163 | 89 | 36707 |
| | 11 | 9 | 13 | 37797 |
| | 56 | 201 | 18 | 38639 |
| | 102 | 13 | 78 | 32617 |
| | 2 | 10 | 2 | 41195 |
| | 48 | 15 | 21 | 40071 |
| | 31 | 10 | 23 | 38624 |
| | 41 | 71 | 73 | 37213 |
| | 504 | 571 | 68 | 38185 |
| | 65 | 542 | 27 | 33213 |
| | 146 | 69 | 4 | 37143 |
| | 9 | 3 | 5 | 39334 |
| | 60 | 9 | 55 | 16340 |
| | 12 | 18 | 10 | 30590 |
| Normal | 0 | 1 | 0 | 39367 |
| | 0 | 17 | 0 | 38751 |
| | 0 | 6 | 0 | 37860 |
| | 0 | 0 | 0 | 39476 |
| | 0 | 7 | 0 | 41761 |
| | 0 | 32 | 0 | 37764 |
| | 0 | 18 | 0 | 40058 |
| | 0 | 20 | 0 | 39648 |
| | 0 | 4 | 0 | 38764 |
| | 0 | 0 | 0 | 38466 |
| | 0 | 8 | 0 | 39960 |
| | 0 | 9 | 0 | 37857 |
| | 0 | 11 | 0 | 32204 |
| | 0 | 6 | 0 | 32325 |

| | | | | |
|---|---|---|---|---|
| | 0 | 69 | 0 | 7772 |
| | 0 | 18 | 0 | 31828 |
| | 0 | 5 | 0 | 30116 |

*Table S9. Performance of the VGGC–A on the ambulatory test set, comprised of epileptic and normal EEGs. Each line represents an EEG of a test patient. At the probability threshold of 0.99, the number of true positives (TP), false positives (FP), false negatives (FN) and true negatives (TN) are shown.*

| | TP | FP | FN | TN |
|---|---|---|---|---|
| Epileptic | 89 | 15 | 27 | 18317 |
| | 156 | 155 | 11 | 40220 |
| | 146 | 3109 | 25 | 34761 |
| | 16 | 6 | 8 | 37800 |
| | 66 | 266 | 8 | 38574 |
| | 122 | 72 | 58 | 32558 |
| | 3 | 32 | 1 | 41173 |
| | 59 | 37 | 10 | 40049 |
| | 43 | 26 | 11 | 38608 |
| | 76 | 168 | 38 | 37116 |
| | 544 | 1094 | 28 | 37662 |
| | 63 | 574 | 29 | 33181 |
| | 145 | 163 | 5 | 37049 |
| | 10 | 7 | 4 | 39330 |
| | 87 | 12 | 28 | 16347 |
| | 15 | 20 | 7 | 30588 |
| Normal | 0 | 4 | 0 | 39364 |
| | 0 | 58 | 0 | 38710 |
| | 0 | 110 | 0 | 37756 |
| | 0 | 4 | 0 | 39472 |
| | 0 | 17 | 0 | 41751 |
| | 0 | 117 | 0 | 37679 |
| | 0 | 984 | 0 | 39092 |
| | 0 | 55 | 0 | 39613 |
| | 0 | 28 | 0 | 38740 |
| | 0 | 4 | 0 | 38462 |
| | 0 | 4 | 0 | 39964 |
| | 0 | 177 | 0 | 37689 |
| | 0 | 39 | 0 | 32176 |
| | 0 | 11 | 0 | 32320 |
| | 0 | 156 | 0 | 7685 |
| | 0 | 24 | 0 | 31822 |
| | 0 | 12 | 0 | 30109 |
